# Lysine Acetylation Reshapes The Downstream Signaling Landscape of Vav1 in Lymphocytes

**DOI:** 10.1101/2020.01.07.897017

**Authors:** Sonia Rodríguez-Fdez, Lucía F. Nevado, L. Francisco Lorenzo-Martín, Xosé R. Bustelo

## Abstract

Vav1 works both as a catalytic Rho GTPase activator and an adaptor molecule. These functions, which are critical for T cell development and antigenic responses, are tyrosine phosphorylation-dependent. However, it is not known whether other posttranslational modifications can modulate the signaling output of this protein. Here, we show that Vav1 becomes acetylated in a stimulation- and SH2 domain-dependent manner. We also demonstrate that the acetylation of four residues located in the catalytic, lysine rich and SH2 domains preferentially buffers the Vav1 adaptor function that favors the stimulation of the nuclear factor of activated T cells. As a result, it modifies the signaling diversification properties normally exhibited by Vav1 in lymphocytes. This new regulatory layer is not shared by Vav family paralogs.

## INTRODUCTION

The Vav family is a group of signal transduction proteins that work both as guanosine nucleotide exchange factors (GEFs) for Rho GTPases and adaptor molecules. This family has single representatives in invertebrates and three members in most vertebrate species (Vav1, Vav2, Vav3)^1,2^. These proteins play critical functions in lymphopoiesis, osteogenesis, cardiovascular homeostasis, neuronal-related processes, and nematode tissue rhythmic behaviors^1–13^ They are also involved in human diseases such as cancer, multiple sclerosis, immune-related deficiencies, and the life cycle of a number of pathogens inside mammalian cell hosts^1,14–19^. Vav1 is the family member that displays a more restricted expression, since is preferentially detected in hematopoietic cells^1, 2^. In line with this, its evolutionary origin parallels the development of the adaptive immune system that took place at the level of jawed fish^2^.

Vav proteins contain a complex array of structural domains that include a calponin-homology (CH) domain, an acidic (Ac) region, the catalytic Dbl-homology (DH) domain, a pleckstrin-homology (PH) region, a C1-type zinc finger (ZF) domain, a lysine rich region (KRR), a proline rich region (PRR), and a SH2 domain flanked by N-(NSH3) and C-terminal (CSH3) SH3 regions (**Fig. 1A**). The noncatalytic domains play pleiotropic roles during signal transduction, contributing to the intramolecular regulation of Vav proteins (e.g., CH, Ac, PH, CSH3), the proper conformation of the central catalytic core (PH, ZF), the phosphorylation step (SH2, CSH3), the stability at the plasma membrane (C1–KR motif), and the engagement of GTPase-independent routes (CH, SH3 domains)^1, 2^. These functions are not mutually exclusive. For example, in the case of Vav1, its CH negatively controls the catalytic activity of the protein in cis and, at the same time, participates as an effector region that triggers the phospholipase Cγ (PLCγ)- and Ca^2+^-dependent stimulation of the nuclear factor of activated T cells (NFAT)^1, 2^. Similarly, the Vav1 CSH3 participates in the optimal phosphorylation of the proteins downstream of the T cell receptor (TCR) as well as in the negative regulation of the Notch1 pathway^20,21^.

**FIGURE 1.**
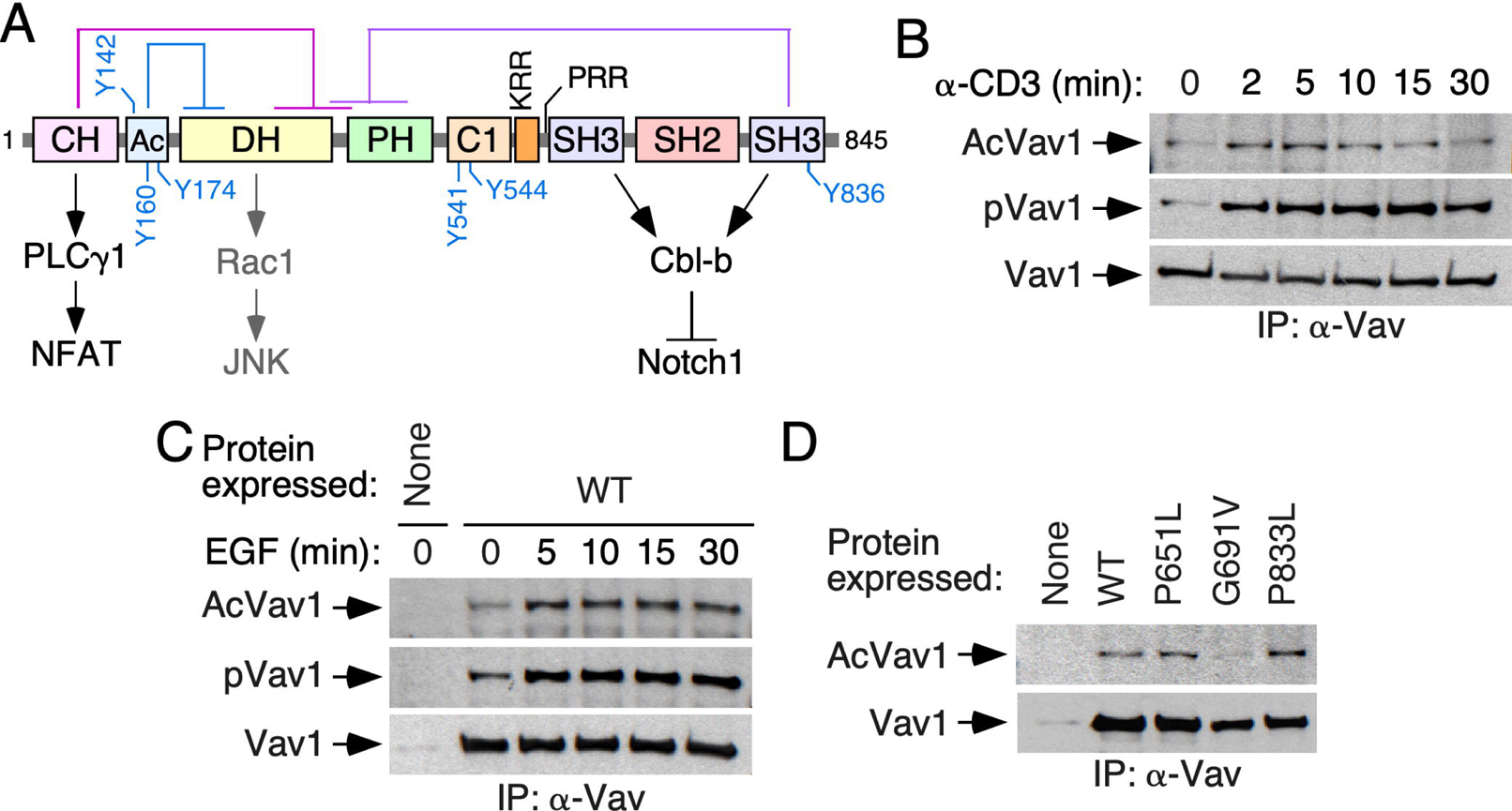
Vav1 lysine acetylation is stimulation- and SH2-dependent. (A) Schematic representation of the structure of Vav1. The intramolecular interactions that maintain the inactive, nonphosphorylated state of the protein are shown on top. The main regulatory phosphosites are shown in blue. The main downstream pathways are shown at the bottom. Abbreviations have been introduced in the main text. **(B)** Kinetics of Vav1 lysine acetylation (top panel) and tyrosine phosphorylation (middle panel) in TCR-stimulated Jurkat cells. The bottom panel shows the amount of endogenous Vav1 immunoprecipitated in each experimental time-point. Similar results were obtained in 6 independent experiments. Ac, acetylated; p, tyrosine-phosphorylated; IP, immunoprecipitation. **(C)** Kinetics of Vav1 lysine acetylation (top panel) and tyrosine phosphorylation (middle panel) in EGF-stimulated COS1 cells. The bottom panel shows the amount of ectopically expressed Vav1 immunoprecipitated in each experimental condition. Similar results were obtained in 3 independent experiments. **(D)** Levels of lysine acetylation (top panel) and tyrosine phosphorylation (middle panel) of the indicated Vav1 mutant proteins ectopically expressed in COS1 cells. The bottom panel shows the amount of immunoprecipitated Vav1 proteins obtained in each case. Similar results were obtained in 3 independent experiments.

The activity of Vav proteins is regulated by tyrosine phosphorylation-dependent conformational changes^1, 2^. In the nonphosphorylated state, Vav proteins adopt a close conformation that is glued together by interactions of both the Vav1 CH–Ac and the CSH3 regions with the central DH and PH domains^22–26^ (**Fig. 1A**). These interactions occlude the effector surfaces of Vav proteins, leading to the inhibition of their catalytic and adaptor-like functions. Upon cell stimulation, the phosphorylation of Vav proteins on specific tyrosine residues (**Fig. 1A**) leads to the release of those autoinhibitory interactions, the exposure of the effectors sites of the molecule, and to full activation^22,23^. Current evidence indicates, however, that the biological activity of Vav proteins can be subjected to additional regulatory layers, including proteolytic cleavage, ubiquitinylation, and arginine methylation^1, 2^. More recently, we have found that the signaling output of activated Vav1 proteins can be modulated by the binding to membrane-resident phosphatidylinostol 5-phosphate (PI5P). This binding, which favors the stability of Vav1 at the plasma membrane, is mediated by the coordinated action of the Vav1 C1 and KR regions^27^. The use of high-throughput mass spectrometry analyses has also revealed that Vav proteins become acetylated on a large number of lysine residues^28–31^. However, the functional impact of this posttranslational modification on the biological activity of these proteins is as yet unknown. Here, we show that lysine acetylation reconfigures the signaling diversification functions of Vav1 in T lymphocytes. Interestingly, this new regulatory layer is not conserved in nonhematopoietic cells and Vav1 paralogs.

## RESULTS

### Acetylation of Vav1 is modulated by upstream stimuli

Given the available proteomics data^28–31^, we investigated whether the acetylation of Vav1 on lysine residues takes place in either a constitutive or a stimulus-dependent manner. To this end, we first used antibodies to acetylated lysine residues to follow the evolution of the acetylation state of the endogenous Vav1 protein in TCR-stimulated Jurkat cells. As shown in **Fig. 1B** (top panel), Vav1 shows some basal acetylation in naïve cells. However, upon TCR stimulation, the protein undergoes a rapid and transient increase in the lysine acetylation levels. These kinetics mimic those found in the case of the phosphorylation of Vav1 on tyrosine residues under the same experimental conditions (**Fig. 1B**, second panel from top). The reblotting of the same blots with antibodies to Vav1 confirmed similar amounts of the immunoprecipitated protein in each of the time-points interrogated in these experiments (**Fig. 1B**, bottom panel). Similar results were observed when the acetylation of ectopically expressed Vav1 was monitored upon stimulation of serum starved COS1 cells with epidermal growth factor (EGF) (**Fig. 1C**). The acetylation of Vav1 is not impaired by inactivating mutations in any of the its two SH3 domains (**Fig. 1D**). By contrast, it is diminished when the Vav1 SH2 is inactivated by a point mutation (G691V) (**Fig. 1D**). These results indicate that, similarly to the tyrosine phosphorylation step, the lysine acetylation of Vav1 is both stimulation- and SH2-dependent.

### Vav1 is acetylated on multiple lysine residues

The key regulatory phosphosites of Vav proteins become phosphorylated upon stimulation of multiple upstream receptors and cell types^1, 2^. To investigate the level of conservation of the Vav1 lysine acetylation landscape across cell types, we next carried out mass spectrometry analyses using immunoprecipitated Vav1 obtained from exponentially growing COS1 cells. Tryptic peptides were considered to harbor acetylated lysine residues when they fulfilled two concurrent criteria: **(i)** Presence of the diagnostic ions with a 126.1 to 143.1 mass to charge ratio^32^. **(ii)** A delay in the retention time when compared to the nonacetylated counterpart. An example of such an identification is shown in **Figure 2A** for the acetylated version of the K^374^ residue. Using this approach, we identified six acetylated residues that were already found in previous high-throughput proteomic experiments (K^194^, K^222^, K^252^, K^587^, K^716^, K^782^) (**Fig. 2B**, residues in green)^28-31^. In addition, we found seven new ones that had not been detected before (K^292^, K^335^, K^353^, K^374^, K^429^, K^775^, K^815^) (**Fig. 2B**, residues in red color). Eight acetylation sites described in previous studies were not found in our analyses (**Fig. 2B**, black residues)^28-31^. All those acetylation sites are surfaced-exposed in the Vav1 3D structure (**Fig. 2C-G**) and cluster within relevant domains of the protein such as the DH Rac1 binding site (**Fig. 2B, C**), the DH surface opposite to the catalytic site (**Fig. 2B, C, D**), the PH (**Fig. 2B, D**), the KRR (**Fig. 2B, E**), the SH2 (**Fig. 2B, F**), and the CSH3 (**Fig. 2B, G**). However, they display in general low levels of phylogenetic conservation in the Vav1 analogs and paralogs that have been identified to date in both invertebrate and vertebrate species (**Fig. 2B** and **S1**)^2^. The comparison with data from previous cell-wide acetylome studies indicate that approx. 28% of the sites identified here are also conserved both in hematopoietic and nonhematopoietic cells (**Fig. 2B**). According to the levels of in vivo acetylation determined in the foregoing mass spectrometry analyses, we found residues that were poorly (less than 1% of the total protein; e.g., K^194^, K^222^, K^252^, K^292^, K^353^, K^587^), mildly (10% of the total protein; e.g., K^335^), and highly (20-35% of the total protein; e.g., K^716^, K^782^) acetylated in vivo (**Fig. 2B**, bottom panel).

**FIGURE 2.**
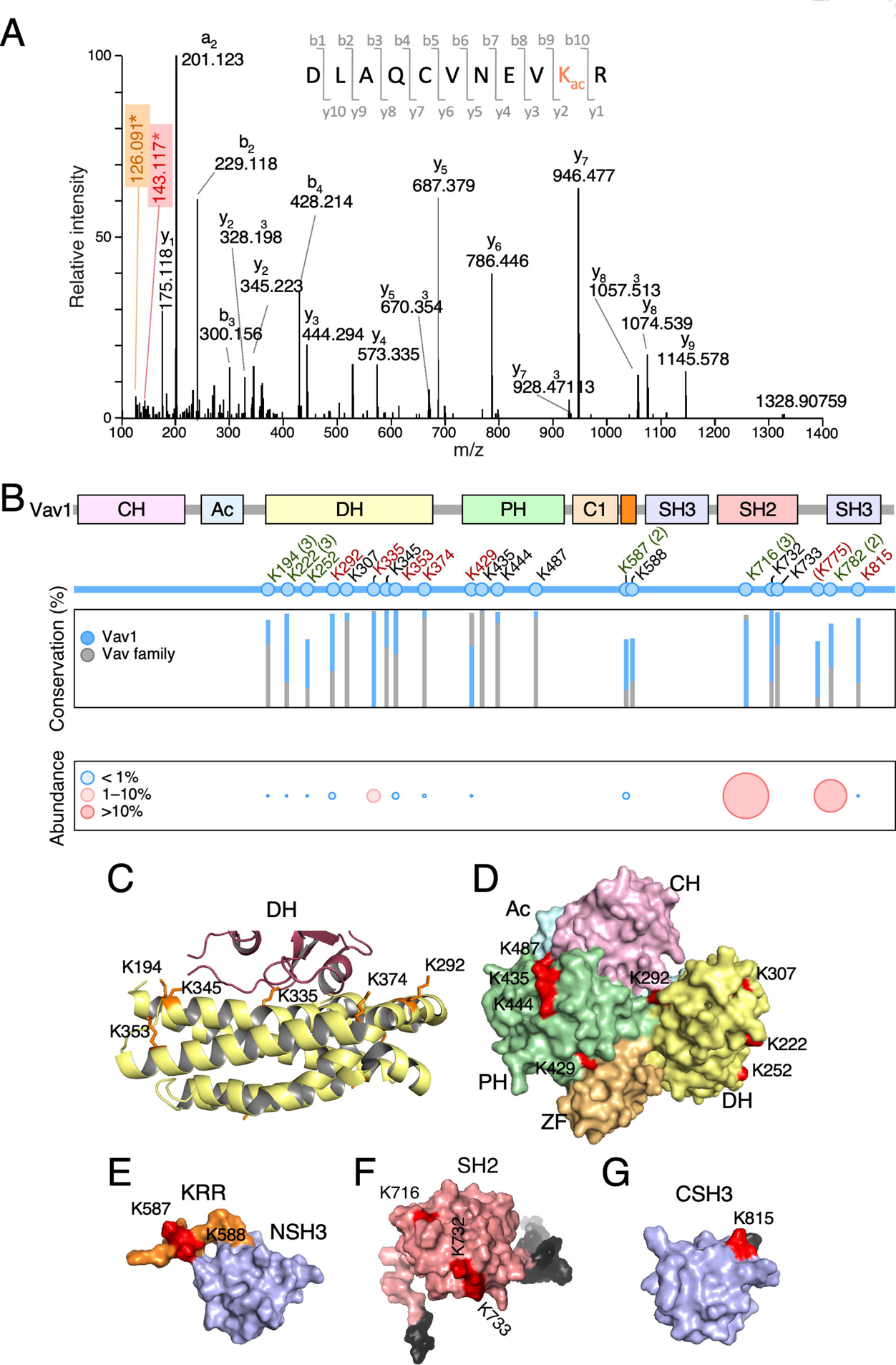
Vav1 is acetylated on multiple lysine residues. (A) Example of the tandem mass spectrometry spectrum of a peptide DLAQCVNEV**K^374^** R containing the acetylated Lys^374^ residue (in bold), showing the acetylated lysine immonium ion at mass-to-charge (m/z) ratio of 143 (red box) and its derivative displaying a m/z ratio of 126 (brown box, generated by loss of ammonia). **(B)** *Top*, main Vav1 lysine acetylation sites identified by mass spectrometry. New acetylation sites found in our study are shown in red. Residues found in our study and previous reports are indicated in green. The number of times in which these acetylation sites have detected before this study is shown in brackets. If only once, the number is not indicated. Previously identified residues that were not found in our analyses are indicated in black. Lys^775^ (in round brackets) is not conserved in mouse Vav1. *Middle*, phylogenetic conservation of the indicated acetylated residues in all Vav1 analogs (blue bars) and paralogs (grey bars) identified to date. *Bottom*, levels of acetylation of indicated residues found in vivo. **(C)** 3D structure of the Vav1 DH domain (yellow) and part of the Rac1 switch regions (brown). The side chains of the indicated lysine residues are shown in orange color. **(D)** 3D structure of the CH–Ac–DH–PH–ZF regions of Vav1 in the inactive conformation. Acetylated residues are shown in red. **(E-G)** 3D structure of the Vav1 KRR (orange) and CSH3 domain (E), the Vav1 SH2 (F), and the Vav1 CSH3 domain of Vav1 (G) with indicated acetylated residues shown in red. In F and G, N- and C-terminal extensions of the domains are indicated in gray. In G, the Lys^782^ residue could not be depicted since it is not included in the available 3D structure.

### The acetylation of the Lys^222^ and Lys^252^ residues modulates Vav1 signaling output

Given the large number of residues involved, we decided to characterize a selection of the Vav1 acetylation sites that fulfilled a number of criteria: **(i)** Detection in more than one mass spectrometry study (K^222^, K^252^, K^587^, K^716^, K^782^; **Fig. 2B**). **(ii)** For those detected in a single study, localization on a potentially important region of the protein (K^335^, K^374^; **Fig. 2B**) or in the neighborhood of one of sites that were found in previous proteomics experiments (K^588^; **Fig. 2B,E**). **(iii)** Level of conservation when compared with Vav1 homologs (**Fig. 2B** and **S1**). To investigate the relevance of those acetylation sites, we mutated each of them into either Arg or Gln. These mutations are usually utilized in protein acetylation studies to mimic the constitutive deacetylated and acetylated states of lysine residues, respectively^28,33,34^. We then tested the biological activity of each of those mutant proteins using luciferase reporter-based c-Jun N-terminal kinase (JNK) and NFAT activity assays in Jurkat cells. Vav1 activates JNK in a catalysis- and Rac1-dependent manner^23,26,35^. Therefore, the activity of this downstream kinase is traditionally used as a surrogate biological readout to measure the activation levels of the Vav1 catalysis-dependent pathways (**Fig. S2A**)^23,35^. Depending on the experiment, we used in these experiments either the wild-type (Vav1^WT^) or a constitutively-active (Vav1^Δ1-186^) version of Vav1. Vav1^Δ1-186^ exhibits tyrosine-phosphorylation activity owing to the lack of the autoinhibitory N-terminal CH and Ac regions (**Fig. S2A**)^26^. As a result, it yields higher levels of activation of the Rac1–JNK pathway than the wild-type counterpart. However, due to the lack of the CH region, this protein cannot activate the NFAT pathway^23,26^ (**Fig. S2A**). The stimulation of the NFAT pathway by Vav1 is mediated by a GTPase-independent, adaptor-like mechanism that involves the PLCγ-mediated stimulation of the nuclear translocation and activation of this transcriptional factor (**Fig. S2A**). The optimal activation of this pathway requires synergistic inputs from TCR-triggered signals (**Fig. S2A**). Due to this, the stimulation of NFAT is boosted by the engagement of the TCR even when using constitutively-active versions of Vav1. Due to this, this assay is conventionally used to monitor the activation levels of this adaptor function of Vav1^36-38^. In this assay, the mutations in the acetylation sites were generated both in the case of the full-length protein and one of its C-terminally truncated versions (Vav1^Δ835-845^). The latter protein displays, similarly to Vav1^Δ1-186^, constitutive, tyrosine phosphorylation-independent biological activity due to the lack of the autoinhibitory CSH3 domain^23^. However, unlike the case of Vav1^Δ1-186^ (Refs^23,26,35^), Vav1^Δ835-845^ is capable of activating the NFAT pathway since it still keeps the N-terminal CH domain^23^. The use of this mutant allowed us to discriminate whether the alterations generated by the acetylation site mutations on Vav1 signaling were related to either the early phosphorylation-dependent Vav1 activation step (in that case, the mutants should affect the biological activity of Vav1^WT^ but not of Vav1^Δ835-845^) or to subsequent effects in the effector signaling phases of the activated protein (in this case, the mutations on the acetylation sites must also affect the signaling output of Vav1^Δ835-845^). Prima facie, we considered that a lysine acetylation site was functionally relevant when it fulfilled two criteria: **(i)** That the Lys to Arg or the Lys to Gln mutant versions of Vav1 on those sites elicited changes in either the NFAT or JNK activity when compared to the WT counterpart. **(ii)** That those effects were opposite in the case of the Lys to Arg and Lys to Gln mutants (e.g., effect/no effect, activation/inactivation, or vice versa). Due to this, we did not consider as bona-fide, functionally relevant those sites in which the Lys to Arg and Lys to Gln mutations elicited the same effect on Vav1 biological activity. It should be noted, however, that this latter criterion might lead to rule out functionally relevant sites if the lysyl group of the mutated lysine residue performs intramolecular roles in the protein that cannot be replaced by the side group of the newly incorporated amino acid residue (see **Discussion**).

Using this experimental approach, we first focused on lysine residues located within the Rac1 binding site of the catalytic DH domain (K^335^, K^374^; **Fig. 2B**, C). When tested in JNK assays, we found that the targeting of any of those two lysine residues leads to a marked reduction in the biological activity of the constitutively-active Vav1^Δ1-186^ (**Fig. S2B,C**). This inhibitory effect is seen in versions of Vav1^Δ1-186^ bearing either Lys to Arg or Lys to Gln mutant versions of these residues (**Fig. S2B,C**). Hence, those two acetylation sites do not fulfill the stringent criteria used in our study to be considered as functionally relevant. When tested in NFAT assays, we did find that the Vav1^K335R^ (**Fig. S2D**, left panel; **Fig. S2E**) and the Vav1^K335Q^ (**Fig. S2F**, left panel; **Fig. S2E**) mutants promote a slight reduction and elevation in the stimulation of this transcriptional factor when compared to Vav1^WT^, respectively. The effect of those mutations is lost when tested in the context of the constitutively-active Vav1^Δ835-845^ protein (**Fig. S2D**, right panel; **Fig. S2E**), suggesting that these effects are not associated with the activation step of the full-length protein. Similar results were obtained when using the two mutant versions of the Lys^374^ residue (**Fig. S2F,G**). Given that the Lys^335^ and Lys^374^ residues are not located within the effector CH domain, these results suggest that the effect derived from the mutation of those sites has to be related to conformational changes in the full-length protein that favor a better exposure of the CH effector interface. A similar effect has been observed before when analyzing mutations targeting key Ac-, ZF-, and CSH3-located phosphosites^23,35^.

Next, we tested the functional role of the acetylation sites (K^222^, K^252^) present in a region of the DH domain opposite to the catalytic site (**Fig. 2B-D**). We observed that the deacetylation-mimicking K222R and K252R mutants promote a very slight, although statistically significant increase in the activity of the full-length Vav1 protein both in nonstimulated and TCR-stimulated cells (**Fig. 3A,B**, left panels). By contrast, the acetylation-mimicking K222Q and K252Q mutants did not elicit any statistically significant change in the activity of the full-length protein. However, when combined together, the double K222Q+K252Q mutant causes a significant reduction in the activity of full-length Vav1 both in nonstimulated and stimulated cells (**Fig. 3A, B**, right panels). This compound mutation, however, does not elicit any change in the activation levels of the JNK by Vav1^Δ1-186^ (**Fig. 3C**). Thus, these two residues do seem to contribute cooperatively to the downregulation of the adaptor function of Vav1 that leads to the stimulation of the transcriptional factor NFAT.

**FIGURE 3.**
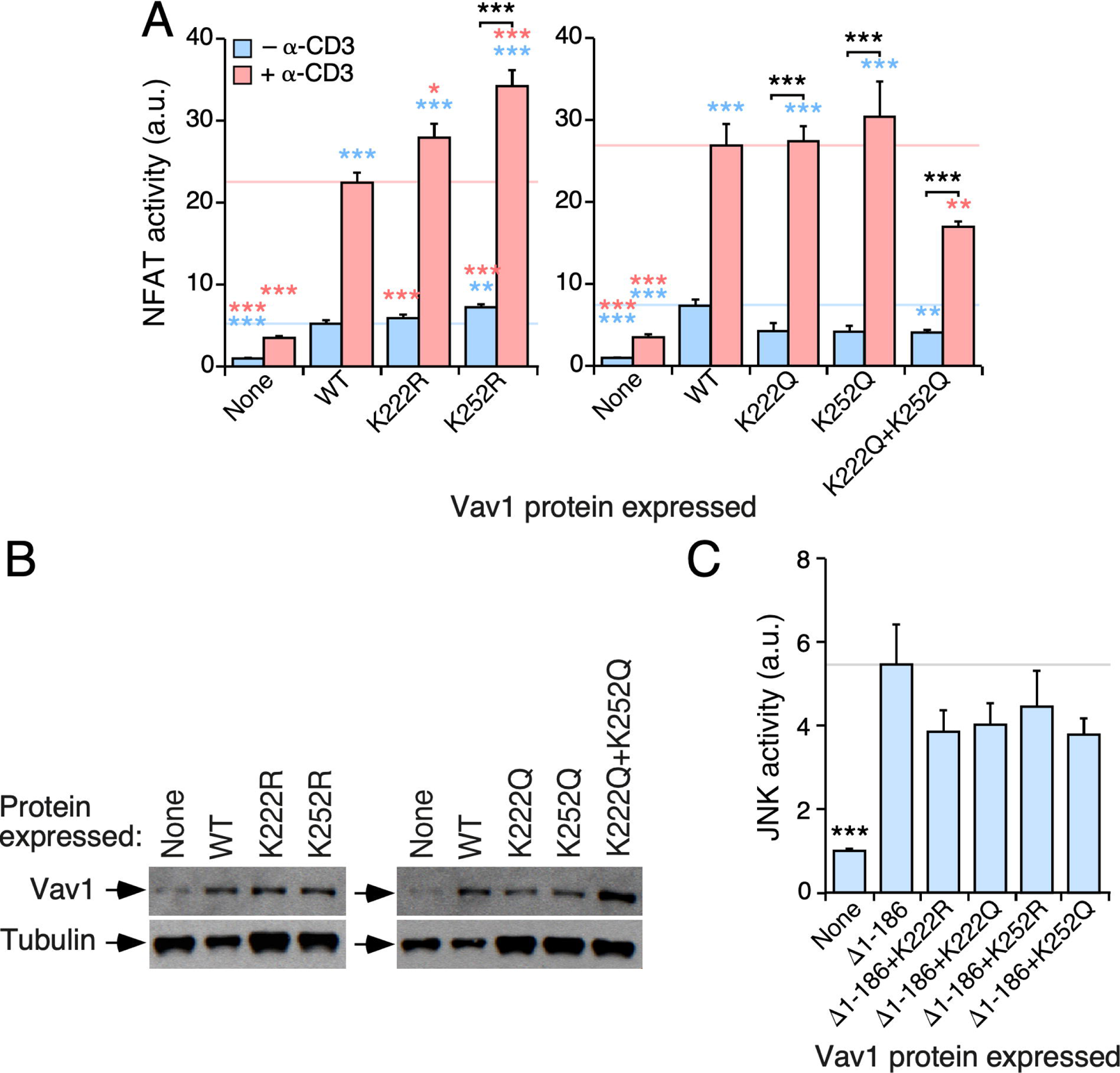
The acetylation residues K^222^ and K^252^ affect the Vav1-mediated stimulation of the downstream NFAT pathway. (A) Activation of NFAT by the indicated ectopically-expressed Vav1 proteins (bottom) in nonstimulated and TCR-stimulated Jurkat cells (inset). Data represent the mean ± SEM. *, *P* < 0.05; **, *P* < 0.01; ***, *P* < 0.001 using the Mann–Whitney U test. Blue and salmon asterisks indicate the significance level compared with nonstimulated and TCR-stimulated Vav1^WT^-expressing cells, respectively. Black asterisks refer to the *P* values obtained between the indicated experimental pairs (in brackets). *n* = 3 independent experiments, each performed in triplicate. **(B)** Representative immunoblots showing the abundance of the ectopic proteins in the experiments shown in A (top panels). Tubulin α was used as loading control (bottom panels). **(C)** Activation of JNK by the indicated ectopically-expressed Vav1 proteins (bottom) in nonstimulated Jurkat cells. Data represent the mean ± SEM. ***, *P* < 0.001 using the Mann–Whitney U test. *n* = 3 independent experiments, each performed in triplicate.

### The acetylation of the KRR-localized Lys^587^ residue affects Vav1 signaling output

We next analyzed the potential contribution of the acetylation of Lys^587^ and Lys^588^ to the biological activity of Vav1. These two residues are potentially interesting, since they are located in the Vav1 KRR that mediates interactions with phosphatidylinositol monophosphates^27^ (**Fig. 2B,E**). We found that the K587Q mutation (that mimics the constitutively acetylated state of this residue), but not the K587R missense change (that mimics the deacetylated state of this site), promotes a marked downregulation of the NFAT activity of the full-length Vav1 protein when tested both in nonstimulated and TCR-stimulated Jurkat cells (**Fig. 4A**, left panel; **Fig. 4B**, two top panels). It also induces a slighter inhibition of full-length Vav1-triggered JNK activity (**Fig. 4A**, right panel, **Fig. 4B**, two bottom panels). The negative effect of the K587Q mutation is eliminated when tested in the context of the Vav1^CAAX^ protein (**Fig. 4C,D**), thus indicating that the defect is related with the stable association of the mutant protein with the plasma membrane. The same behavior was observed before in the case of mutants targeting other residues of the Vav1 KRR^27^. The activity of Vav1^K588R^ and Vav1^K588Q^ proteins show no statistically significant alteration when tested in both NFAT and JNK assays (**Fig. 4E,F**). This result indicates that the Lys^587^ residue is a functionally relevant Vav1 acetylation site.

**FIGURE 4.**
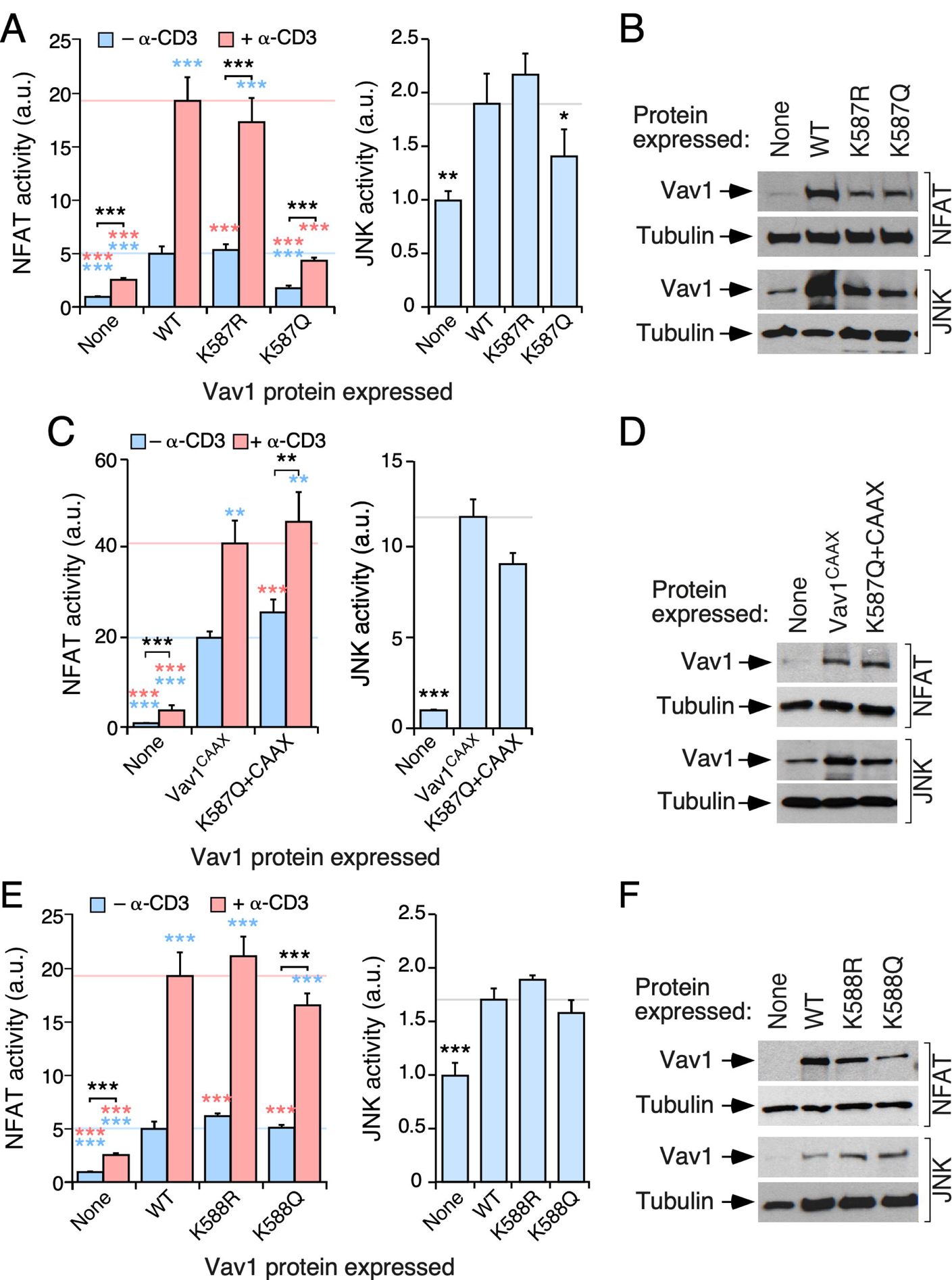
The acetylation residue Lys^587^ is important for Vav1 signaling output. **(A,C,E)** Activation of NFAT (A,C,E; left panels) and JNK (A,C,E; right panels) by indicated Vav1 proteins in nonstimulated and TCR-stimulated Jurkat cells. Data and *P* values are depicted as in Figure 3A (*n* = 3 independent experiments, each performed in triplicate). **(B,D,F**) Representative immunoblots showing the abundance of the ectopic Vav1 proteins in the experiments shown in A (B), C (D), and E (F). Tubulin α was used as loading control in all cases.

### The acetylation of the SH2-located Lys^716^ residue limits Vav1 signaling output

This residue is prima facie appealing because: **(i)** It shows high levels of acetylation in cells according to our mass spectrometry determinations (**Fig. 2B**). **(ii)** It is the only acetylation site conserved in all mammalian Vav family proteins (**Fig. 2B** and **S1**). **(iii)** It is located in the vicinity of the grove of the Vav1 SH2 domain that participates in the interaction with phosphotyrosine peptides (**Fig. 2F**). The mutations of this residue promote an increase (in the case of the deacetylation-mimicking Lys to Arg mutation) and a reduction (in the case of the acetylation-mimicking Lys to Gln mutation) on Vav1-triggered NFAT activity in both nonstimulated and TCR-stimulated cells (**Fig. 5A,B**). The same effect is observed when the K716Q mutations are analyzed in the context of the hyperactive, phosphorylation-independent Vav1^Δ835-845^ version (**Fig. 5A**, right panel; **Fig. 5B**, two bottom panels). By contrast, the K716R and the K716Q versions of full-length Vav1 (**Fig. 5C,D**; left panels) and Vav1^Δ1-186^ (**Fig. 5C,D**; right panels) exhibit normal levels of JNK activation when compared to the nonmutated counterparts. Thus, similarly to the rest of mutations tested so far, the acetylation of this residue seems to primarily affect the adaptor-like rather than the catalysis-based downstream activities of Vav1.

**FIGURE 5.**
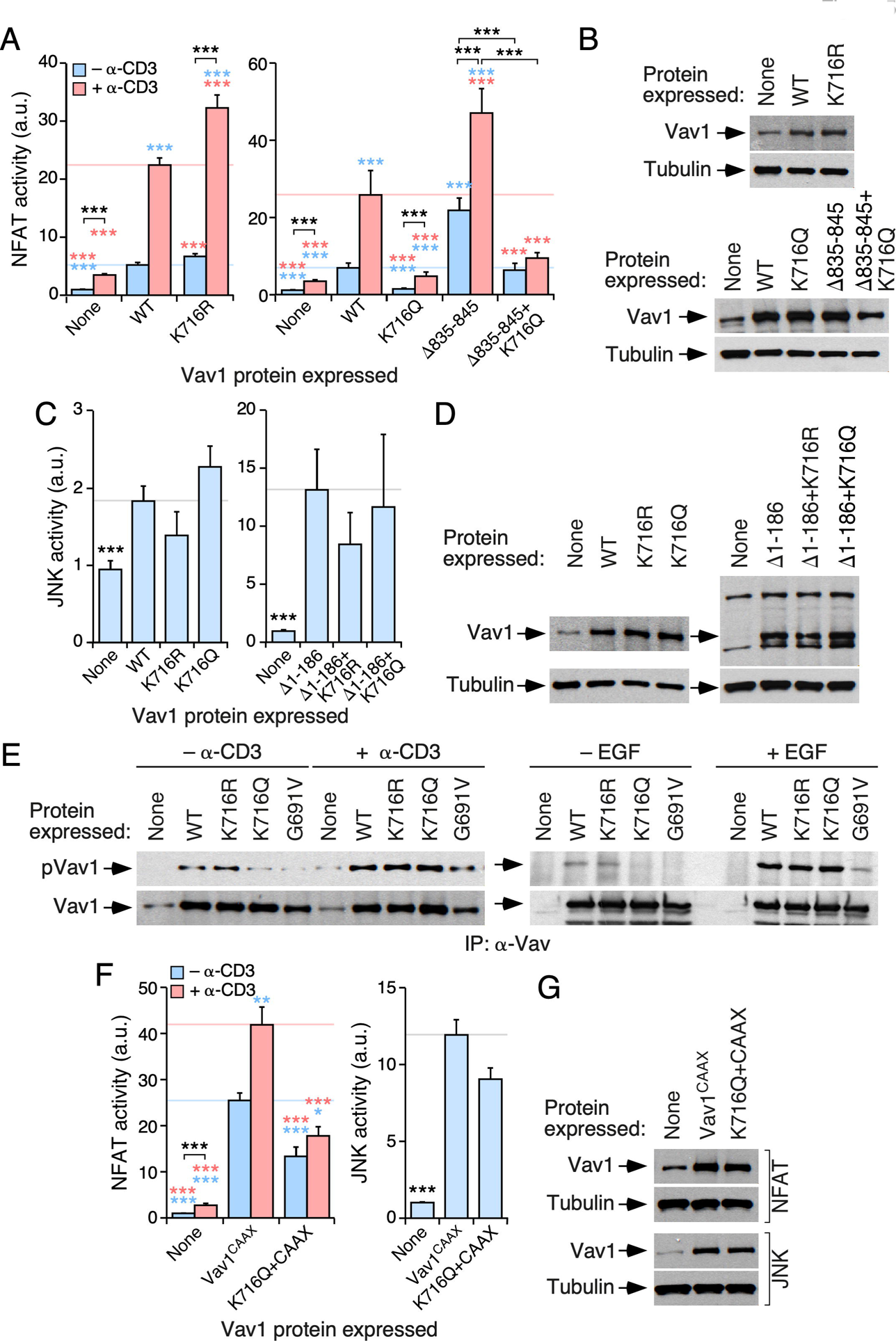
The acetylation residue K^716^ limits Vav1 signaling output. (A,C) Activation of NFAT (A) and JNK (C) by the indicated Vav1 proteins in nonstimulated and TCR-stimulated Jurkat cells. Data and *P* values are depicted as in **Figure 3A** (*n* = 3 independent experiments, each performed in triplicate). **(B,D)** Representative immunoblots showing the abundance of the indicated Vav1 proteins in the experiments shown in A (B) and C (D). Tubulin α was used as loading control in all cases. **(E)** Levels of tyrosine phosphorylation of the indicated Vav1 proteins immunoprecipitated from nonstimulated and stimulated Jurkat (top panels on the left) and COS1 (top panels on the right) cells. The stimulation conditions are indicated on top. The bottom panels show the amount of endogenous Vav1 immunoprecipitated in each sample. Similar results were obtained in 3 independent experiments. **(F)** Activation of NFAT (left panel) and JNK (right panel) by indicated Vav1 proteins in Jurkat cells under nonstimulation and TCR-mediated stimulation conditions. Data and *P* values are depicted as in Figure 3A (*n* = 3 independent experiments, each performed in triplicate). **(G)** Representative immunoblots showing the abundance of the ectopic Vav1 proteins in the transfected cells used in F. Endogenous tubulin α has been used as loading control.

Given the importance of the SH2 domain for the tyrosine phosphorylation of Vav1^1,2,39^, we next investigated the possible impact of the Lys^716^ mutations on the phosphorylation of the protein in both Jurkat and COS1 cells. We found that the K716Q mutation, but not the K716R one, promotes a marked reduction in the tyrosine phosphorylation levels of full-length Vav1 in nonstimulated Jurkat (**Fig. 5E**, left panel) and serum starved COS1 (**Fig. 5E**, right panel) cells. This effect is similar to that found when using the G691V mutation that inactivates the phosphotyrosine binding activity of the Vav1 SH2 domain (**Fig. 5E**). However, unlike the case of the G691V missense change, the full-length Vav1 protein bearing the K716Q mutation exhibits levels of tyrosine phosphorylation comparable to those found in Vav1^WT^ when monitored in both TCR-stimulated Jurkat (**Fig. 5F**, left panel) and EGF-stimulated COS1 (**Fig. 5F**, right panel) cells. These results suggest that this mutation reduces, but does not abrogate, the affinity of the Vav1 SH2 domain for its tyrosine phosphorylated ligands. It is unlikely, however, that this is the underlying cause of the deleterious effects of the K716Q mutation on Vav1 activity because: **(i)** The K716Q mutation does not impair the stimulation of JNK by the full-length Vav1 protein **(Fig. 5C,D**), an activity that is also tyrosine phosphorylation-dependent^23,40^. **(ii)** This mutation also impairs the stimulation of NFAT by the phosphorylation-independent Vav1^Δ835-845^ protein (**Fig. 5A,B**). **(iii)** The reduced activity of the Vav1^K716Q^ mutant in NFAT assays cannot be restored by the attachment to the protein of the membrane anchoring signal of H-Ras^41^(**Fig. 5F,G**). This signal promotes high levels of Vav1 activity due to enhanced membrane localization and phosphorylation^26^.

Given that this acetylation site is conserved in both Vav2 (Lys^718^ residue) and Vav3 (Lys^717^ residue) (**Fig. S1**), we next investigated whether the acetylation of this residue could influence the activity of other family proteins. We focused our attention on Vav3, given that Vav2 cannot stimulate the NFAT pathway in Jurkat cells^42^. We could only find minor effects of the K717Q mutation on the biological activity of full-length Vav3 in both nonstimulated and TCR-stimulated cells (**Fig. S3A,B**). Moreover, the K717R mutation does not alter the signaling output of Vav3 (**Fig. S3A,B**). Interestingly, we found that the impact of each of those mutations on the overall tyrosine phosphorylation levels of Vav3 and Vav1 are quite different. Thus, unlike the case of Vav1, we could not find any significant variation in the levels of phosphorylation between Vav3^WT^ and the Vav3^K717Q^ protein. Furthermore, the Vav3^K717R^ mutant unexpectedly exhibited much higher levels of phosphorylation than Vav3^WT^ (**Fig. S3C**) (see above; **Fig. 5E**, left panel). These results indicate that the K^716^ acetylation residue is only functionally relevant in the case of Vav1.

Given the differential effect induced by the attachment of the H-Ras membrane anchoring signal to the Vav1^K587Q^ (rescue of activity) and Vav1^K716Q^ (no rescue) mutants, we decided to investigate whether the enforced membrane localization of Vav1 could restore normal NFAT activity levels in the case of the mutant Vav1^K222Q+K252Q^ protein (**Fig. 3A)**. We found that the Vav1^K222Q+K252Q+CAAX^ version induces levels of NFAT activity similar to those elicited by the control Vav1^CAAX^ protein (**Fig. S3D,E**). These results indicate that the regulation mediated by the Lys^716^ must be mechanistically different from the mode of action of the acetylation sites located in the Vav1 DH and KRR regions.

### Cooperativity of DH and SH2 acetylation sites on Vav1 signaling output

To assess the consequences of the effect of the concurrent elimination of the acetylation Lys^222^, Lys^252^, and Lys^716^ sites on Vav1 activity, we generated versions of the full-length protein bearing triple Lys to Arg (K3xR) and Lys to Gln (K3xQ) mutations in those sites. We found that the ectopically expressed Vav1^K3xR^ mutant displays higher levels of NFAT activity in both nonstimulated and stimulated cells when compared with the wild-type counterpart (**Fig. 6A**, left panel). Conversely, the Vav1^K3xQ^ mutant exhibited a total lack of NFAT activity in the same assays (**Fig. 6A**, right panel). By contrast, we found no statistically significant effects of any of those mutations in Vav1-mediated JNK activity (**Fig. 6B**). Western blot experiments confirmed the proper expression of the ectopic proteins used in these experiments (**Fig. 6C,D**). Further experiments indicated that the K716Q mutation, but not the compound K222Q+K252Q mutation, affects the basal tyrosine phosphorylation state of full-length Vav1 found in nonstimulated Jurkat (**Fig. 6E**, top panels) and COS1 (**Fig. 6E**, bottom panels) cells. Taken together, these results confirm that the acetylation of these three acetylation sites leads to the negative regulation of the Vav1 signaling branch that favors the stimulation of NFAT. Given that there is not an additive effect of the two mutant subsets on Vav1 phosphorylation levels, these results also indicate that the mode of action of the DH and SH2 acetylation sites is mechanistically different.

**FIGURE 6.**
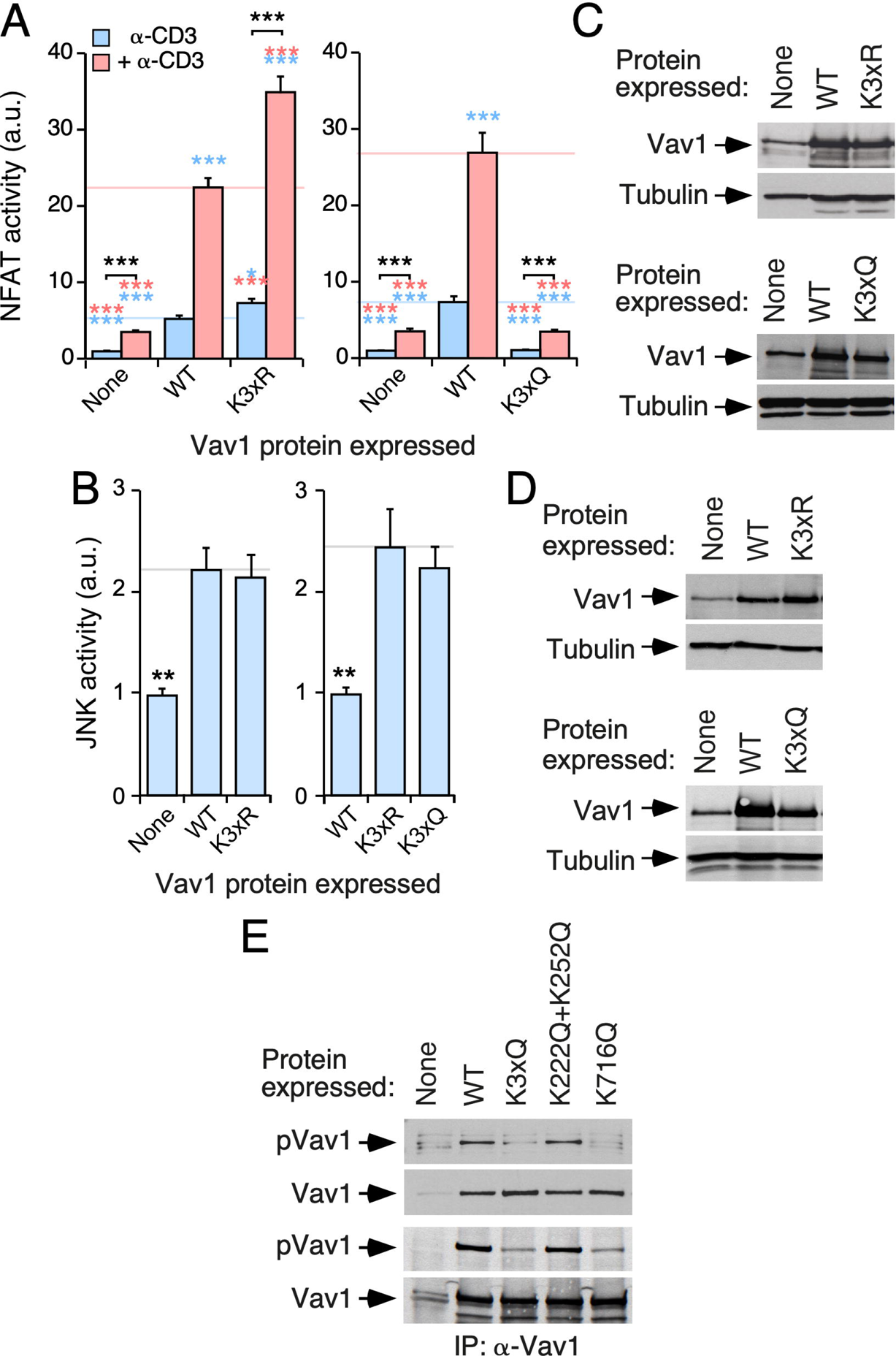
DH-and SH2-located acetylation sites cooperate in the downmodulation of Vav1 signaling output. (A,B) Activation of NFAT (A) and JNK (B) by indicated Vav1 proteins in nonstimulated (A,B) and TCR-stimulated (A) Jurkat cells. Data and *P* values are depicted as in Figure 3A (*n* = 3 independent experiments, each performed in triplicate). **(C,D)** Representative immunoblots showing the abundance of the ectopic Vav1 proteins in the transfected cells used in panels A and B, respectively. Endogenous tubulin α was used as loading control. **(E)** Levels of tyrosine phosphorylation of the indicated Vav1 proteins immunoprecipitated from nonstimulated COS1 cells (top panel). The bottom panel shows the amount of Vav1 immunoprecipitated in each sample. Similar results were obtained in 3 independent experiments.

### Characterization of the acetylation Lys^782^ site present in the Vav1 CSH3 domain

Finally, we analyzed the potential contribution of the Lys^782^ residue to the biological activity of Vav1. Although relatively poorly conserved from a phylogenetic point of view (**Figs. 2B** and **S1**), this site is the second mostly acetylated (22% of the total protein) according to our mass spectrometry analyses (**Fig. 2B**, bottom panel). In this case, we found that the K782R and K782Q mutations promote the same stimulatory effect on the biological activity of full-length Vav1 when tested both in NFAT and JNK assays in Jurkat cells (**Fig. S4A,B**). Given that Lys^782^ is localized in the vicinity of one of the interfaces of the CSH3 domain that contributes to the intramolecular inhibition of the protein^23^, it is likely that the upregulated activity of Vav1^K782R^ and Vav1^K782Q^ proteins is owing to the release of the CSH3-mediated intramolecular inhibition of the protein (see **Discussion**).

## DISCUSSION

In this work, we have shown that Vav1 becomes acetylated on multiple lysine residues in a cell stimulation- and a SH2-dependent manner. Furthermore, our cell-based studies have shown that the acetylation of four lysine acetylation sites (K^222^, K^252^, K^587^, K^716^) contributes to the negative regulation of the biological activity of Vav1 (**Fig. 7**). These acetylation sites can be assigned to two overlapping, although mechanistically different subsets. One of those subsets, which is composed of the lysine residues located in the KRR (Lys^587^) and in the noncatalytic side of the DH domain (Lys^222^, Lys^252^), is involved in the regulation of the stability of Vav1 at the plasma membrane. Consistent with this, we can bypass the negative effect elicited by the acetylation-mimicking mutations on those residues by enforcing the constitutive localization of the mutant protein at the plasma membrane. It is likely that the acetylation of these residues will reduce the interaction of Vav1 with phospholipids residing at the plasma membrane. This idea is in agreement with the recently described role of the Vav1 C1–KRR cassette in the establishment of interactions with membrane-resident phosphatidylinositol monophosphates^27^. The second functional subset includes the Lys^716^ residue. The negative effect derived from the acetylation-mimetic mutation on this residue cannot be overcome by the forced tethering of the mutant Vav1 protein to the plasma membrane, thus suggesting that the acetylation of this site must contribute to the regulation of the biological activity of Vav1 using a different mechanism from the rest of acetylation sites interrogated in this study. Given that the Lys^716^ is located in the phosphopeptide-binding grove of the Vav1 SH2 domain (**Fig. 2B,F**), one possibility is that the acetylation step could reduce the affinity towards the upstream protein tyrosine kinases that phosphorylate Vav1 upon the engagement of the TCR. However, although we did find reduced tyrosine phosphorylation levels of the Vav1^K716Q^ mutant protein in nonstimulated cells, our results are not consistent with that hypothesis since normal Vav1 phosphorylation levels have been found in TCR-stimulated cells. Supporting this interpretation, we have observed that the K716Q mutation also reduces the biological activity of Vav1 mutant proteins that display phosphorylation-independent activity (e.g., Vav1^Δ835-845^)^23^. Those data suggest that the nonacetylated version of this residue could be important to maintain a conformation of the activated protein compatible with optimal downstream signaling. Alternatively, it might contribute to the high affinity binding of a hitherto unknown SH2-binding phosphoprotein that could be important for the engagement of specific signaling branches of Vav1. We currently favor the former possibility, because similar conformational-associated effects on Vav1 activity have been observed when using mutant proteins for specific regulatory phosphosites^23,35^. In any case, further studies will be needed to fully understand whether the effects of the K716Q mutation on Vav1 activity are either *in cis* or *in trans*. Regardless of its specific regulatory function, the Lys^716^ residue is probably one of the most relevant for the regulation of Vav1 activity given its high levels of acetylation found in cells (**Fig. 2B**).

**FIGURE 7.**
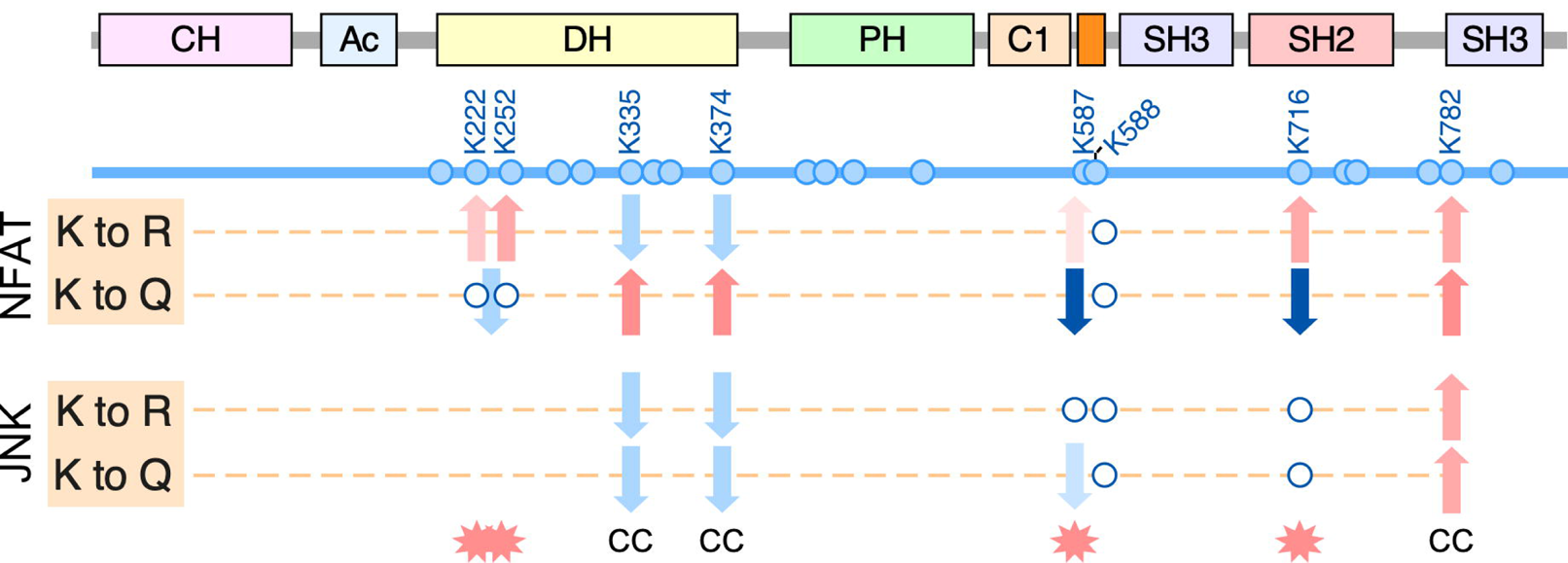
Summary of the results obtained in our work with the indicated deacetylation- and acetylation-mimetic Vav1 mutants. The residues selected for our analyses are indicated in blue font. The biological assays and mutant used are indicated on the left. Red and blue arrows indicate increased and decreased stimulation of the indicated pathway, respectively, when compared to the activity of Vav1^WT^. Empty circles indicate no statistically significant differences in activity when compared to Vav1^WT^. When using a double mutant protein, the arrow is placed between the two circles. Acetylation-dependent regulatory sites according to our experimental criteria are indicated as red stars (bottom). CC, Sites that when mutated modify the Vav1 signaling output via a putative conformational change of the protein (see **Discussion**).

The acetylation-mimetic mutations have a much stronger negative impact on the adaptor-like functions of Vav1. This NFAT-specific effect cannot be attributed to membrane localization issues, since the stimulation of JNK pathway requires the subcellular localization of both Vav1 and Rac1 at the plasma membrane. One plausible hypothesis to explain the signaling branch-specific effects of those mutations is that the optimal stimulation of NFAT would require longer periods of localization of Vav1 at the plasma membrane than the catalysis-dependent pathways, a difference that probably stems from the fact that the former pathway needs the stepwise engagement of a much larger number of signaling elements than the latter one^1,37^. Alternatively, it is possible that the interaction with specific membrane regions (e.g., PI5P enriched) could favor the acquisition of a specific orientation and/or structural conformation by the nonacetylated Vav1 that could facilitate the subsequent interaction with proximal effectors specifically involved in the stimulation of the NFAT pathway. In agreement with this latter possibility, we have shown before that the specific engagement of this pathway requires structural arrangements of the Vav1 molecule more complex than those needed to stimulate the Rac1–JNK axis^35^.

Our results also indicate that the effect of the acetylation-mimetic mutants is stronger than those elicited by the deacetylation-mimetic ones. This is probably a reflection of the low levels of acetylation found for the interrogated residues in cells (**Fig. 2B**). It is also likely that, in order to get a full impact on Vav1 activity, the protein would have to become acetylated at several sites at the same time. This is in agreement with the stronger effect seen with proteins bearing compound mutations in some of those sites. This is similar to the case of the phosphorylation-mediated regulation of Vav1 protein, which also requires the concurrent phosphorylation of several tyrosine residues to become fully biologically competent^23,35^.

The characterization of other acetylation sites has yielded inconclusive results. In the case of Lys^335^ and Lys^374^, this was due to the fact that the acetylation and deacetylation-mimicking mutants elicits similar inhibitory effects on the Rac1–JNK pathway (**Fig. 7**). In the case of the CHS3-located Lys^782^ residue, this was caused by the similar stimulatory effect of the Lys to Arg and Lys to Gln mutations on Vav1 adaptor-like and catalysis-dependent functions (**Fig. 7**). Due to this, we cannot formally prove that the changes in activity of the mutant proteins reflect an intrinsic effect triggered by the acetylation state of these lysine residues. It is worth noting, however, that the similarity in the signaling response induced by antagonistic mutations does not exclude per se that those sites could be relevant from a functional point of view. For example, we would obtain similar biological effects by these antagonistic mutations in the case that the lysyl group of the nonacetylated residue contributes to the formation of key intramolecular interactions within the protein. If that were the case, it is unlikely that the structural function of the lysyl group could be fully recapitulated by the side group of the new amino acid that has been introduced in that position upon the mutagenesis step. Such an scenario has already been seen when using dephosphorylation-(Tyr to Phe) and phosphorylation-mimicking (Tyr to Glu) mutations targeting key Vav1 regulatory phosphosites^22,23,26,40^. For example, in the case of the mutations targeting the Y^174^ site, this effect is due to the fact that the hydroxyl group of the nonphosphorylated version of this residue forms interdomain bonds that are important to maintain the inactive, “close” conformation of the molecule^22^. Hence, it is possible that a similar scenario could apply here in the case of the Lys^335^, Lys^374^ and Lys^782^ residues. If that were the case, the acetylation of the Lys^335^ and Lys^374^ residues could cooperate with the acetylation of the K^222^, K^252^, K^587^, K^716^ residues to reduce both the adaptor-like and catalysis-dependent signaling of the protein. Conversely, the acetylation of the Lys^782^ residue could act in a concerted manner with the phosphorylation of the regulatory tyrosine residues to eliminate the intramolecular inhibitory effect of the CSH3. The functional role of this site would be therefore opposite to the rest of acetylation sites analyzed in this work. Bivalent functions of acetylation sites in the same molecule have been described before^43^. The acetylation residues of Vav1 show low level of phylogenetic (**Fig. S1**) and functional (**Fig. S3**) conservation among Vav family proteins. The KRR-regulatory mechanism is also exclusive for Vav1^2,27^. Thus, unlike the case of the phosphorylation-dependent regulation^2^, these two new mechanisms seem to have been developed during evolution to fulfill Vav1- and lymphocyte-specific regulatory needs.

Although initially associated with nuclear-related functions, it is now known that lysine acetylation plays more widespread roles in cytosolic, plasma membrane, and cellular organelle proteins^43^. In the case of Rho-related pathways, acetylation has been shown to affect the activity of a variety of Rho GDP dissociation inhibitors, guanosine nucleotide exchange factors (e.g., Tiam1, Net1), cytoskeletal related proteins (e.g., cortactin), and microtubule components^43-47^. Our data have identified Vav1 as a new member of this poorly characterized club of acetylation-regulated signaling and cytoskeletal proteins. They also indicate that, in addition to the regulation of the on/off state previously described in other proteins, this posttranslational modification contributes to reshaping the signaling diversification landscape triggered by Vav1 in antigen-stimulated T cells.

## MATERIALS AND METHODS

### Mammalian expression vectors

All the Vav family constructs used in this work encode versions of the murine species and were DNA sequence-verified in our Genomics Facility. Plasmids encoding Vav1^WT^ (pJLZ52), Vav1^Δ^(pMJC10), Vav1^Δ^(pSRF49), Vav1 (pSRF46), Vav1 (pSRF93), Vav1^G691V+CAAX^ (pSRF96), Vav1^Y174E^ (pMB123), and EGFP-Vav1^WT^ (pSRM3) were previously described^23,26,35,48^. The pNFAT–Luc (Cat. number 17870) plasmid was obtained from Addgene, the pSRE–luc (Cat. number 219081), the pFR–Luc and pFA2– cJun plasmids from Stratagene (now, Agilent Technologies), and the pRL–SV40 plasmid from Promega. The rest of the plasmids encoding Vav1 mutant proteins were generated in this work by site-directed mutagenesis using the high-fidelity NZYProof DNA polymerase (Cat. #14601, NZYTech) and the appropriate combination of mutation-bearing oligonucleotides (**Table S1**).

### Immunological reagents

The rabbit polyclonal antibodies to the Vav1 DH domain (Lab reference number 302-5), phospho-Y^174^ (Lab reference number 613), phosphoY^280^ (Lab reference number 595) and phosphoY^836^ (Lab reference number 622) have been described elsewhere^23,35,40^. Other antibodies used in this study include those specific to human CD3 (UCHT1 clone, Cat. number 217570, Merk-Millipore), Vav1 (Cat. number sc-8039, Santa Cruz Biotechnology), phosphotyrosine (Cat. number sc-7020, Santa Cruz Biotechnology), acetyl-lysine (Cat. number AB3879, Chemicon), CD98 (Cat. number ab108300, Abcam), GAPDH (Cat. number sc-25778, Santa Cruz Biotechnology), polyhistidine (Cat. number H-1029; Sigma), HA (Cat. number 5017; Cell Signaling) and tubulin α (Cat. number CP06, Calbiochem). Rhodamine-labeled phalloidin (Cat. number PHDR1) was from Cytoskeleton.

### Cell culture

Jurkat cells were obtained from the ATCC and grown in RPMI-1640 medium supplemented with 10% fetal calf serum, 1% L-glutamine, penicillin (10 μg/mL) and streptomycin (100 μg/mL). COS1 cells were obtained from the ATCC and grown in DMEM supplemented with 10% fetal calf serum, 1% L-glutamine, penicillin (10 μg/mL) and streptomycin (100 μg/mL). All tissue culture reagents were obtained from Gibco. All cell lines were maintained at 37°C in a humidified, 5% CO_2_ atmosphere. When required, cells were stimulated for the indicated periods of time with antibodies to human CD3 (7.5 μg/mL, Cat. number 217570, Merck; Jurkat cells) and EGF (50 ng/mL, Cat. number E9644, Sigma; COS1 cells). In the latter case, cells were previously starved without serum for 3 hours.

### Western blotting

Immunoblot analyses were carried out as described elsewhere^35^.

### Immunoprecipitations

In the case of COS1 cells, we ectopically expressed the indicated proteins using the diethylaminoethyl-dextran (10 mg/mL, Cat. #D9885, Sigma)/chloroquine (2.5 mM, Cat. #C6628, Sigma) method^49^. Two 10-cm diameter plates were transfected per condition using 10 μg of plasmid per plate. 48 hours post-transfection, cells were washed in a phosphate-buffered saline solution and lysed with the aid of a scrapper in 1 mL of RIPA buffer [10□mM Tris-HCl (pH□7.5), 150□mM NaCl, 1% Triton X-100 (Cat. number X100, Sigma), 1 mM Na_3_VO_4_ (Cat. number S6508, Sigma), 1 mM NaF (Cat. number S7920, Sigma) and the CØmplete protease inhibitor cocktail (Cat. number 11836145001, Roche)]. Cellular extracts were kept 5 min on ice and subsequently centrifuged at 14 000 rpm for 10 min at 4 °C to eliminate cell debri. The supernatants were then incubated with either 7 μL of an antibody to Vav1 or 5 μL of an antibody to acetylated lysine residues overnight at 4°C. Immunocomplexes were then collected with Gammabind G-Sepharose beads (Cat. number GE17-0885-01, GE Healthcare), washed three times in RIPA buffer, resuspended in SDS-PAGE buffer, boiled for 5 min, electrophoresed, and subjected to immunoblot analyses with the indicated antibodies.

In the case of Jurkat, 10 × 10^6^ cells were transfected with 20μg of DNA per condition using the Program #5 of the Neon transfection system (Cat. number MPK5000, Invitrogen) according to the manufacturer’s instructions. After 48 hours, the cells were washed, transferred to a 1.5 mL Eppendorf tube and, after low speed centrifugation at 4 °C, resuspended in lysis buffer with extensive vortexing. The cell extracts obtained were processed as above. However, in the case of immunoprecipitation experiments using GFP-Vav1, the cell lysates were incubated with the GFP-Trap reagent (Cat. number gta-100, ChromoTek) for 1 hour at 4 °C. Washed immunocomplexes were collected by centrifugation and subjected to immunoblot analyses as above. In all cases, total cellular lysates were analyzed in parallel to monitor the expression of the ectopically expressed proteins used in each experiment.

### Mass spectrometry analyses

Upon the GFP-trap-mediated immunoprecipitation, the GFP-Vav1 proteins were separated by one-dimensional SDS-PAGE electrophoresis and Coomassie blue-stained. The area of the gel containing the GFP-Vav1 (according to molecular weight mobility) was excised and subjected to in-gel digestion with trypsin following a modified version of a protocol described elsewhere^50^. To this end, the gel pieces were destained at 37°C for 15 min using a solution of 50% acetonitrile in 50 mM sodium bicarbonate. Subsequently, the protein reduction and alkylation steps were performed using 10 mM DTT (56 °C, 45 min) and 55 mM iodoacetamide (room temperature, 30 min). Upon digestion with trypsin (6.25 ng/mL) at 37 °C for 18 hours, the peptide-containing solutions were acidified with formic acid (FA) and desalted by using C18-Stage-Tips columns^51^, partially dried (how), and stored at −20°C. For mass spectrometry analysis, the peptides were dissolved in 0.5% FA/3% acetonitrile (ACN), loaded onto a trapping column (nanoACQUITY UPLC 2G-V/M Trap Symmetry 5 μm particle size, 180 μm × 20 mm C18 column; Cat. number 186006527, Waters Corp., Milford/MA, USA), and separated using a nanoACQUITY UPLC BEH C18 column (1.7 μm, 130 Å and a 75 μm × 250 mm; Cat. number 186003545, Waters Corp., Milford/MA, USA) at 40°C and with a linear gradient ranging from 7 to 35% of solvent B (ACN/0.1% FA; flow rate: 300 nL/min over 30 min and 5 min to 55%). The LTQ-Orbitrap Velos was operated in the positive ion mode applying a data-dependent automatic switch between survey mass spectra (MS) scan and tandem mass spectra (MS/MS) acquisition. MS scans were acquired in the mass range of m/z 400 to 1400 with 30,000 resolution at m/z 400, with lock mass option enabled for the 445.120025 ion^52^. The 10 most intense peaks having ≥2 charge state and above 500 intensity threshold were selected for fragmentation by high energy collision-induced dissociation (HCD) at 42% normalized energy, 100 ms activation time and 2 m/z precursor isolation width. Maximum injection time was 1,000 ms and 500 ms for survey and MS/MS scans, respectively. Automatic gain control was 1 × 106 for MS and 5 × 104 for MS/MS scans. Dynamic exclusion was enabled for 90 s. MS/MS spectra were acquired in the Orbitrap with 7,500 resolution. A parent mass list with m/z corresponding to possible acetylated peptides was build using Skyline and included in the acquisition method^53^.

Mass spectra were analyzed using the SEQUEST HT algorithm of the Proteome Discoverer software (Cat. number OPTON-30795, Thermo Fisher Scientific). All tandem mass spectra were searched against a custom database containing Uniprot complete mouse sequences and Mann contaminants. The search parameters were fully-tryptic digestion with a maximum of 2 missed cleavages, 20 ppm and 0.02 Da mass tolerances for precursor and product ions respectively, carbamidomethylated cysteines, variable oxidation of methionine and variable acetylation on lysine. 1% false discovery rate using Percolator^51^ was used for peptide validation. In order to evaluate the proportion of acetylated peptides, a mass deviation of 2 ppm was used for precursor ion area detection, and peptides with 5% false discovery rate were also included in the calculation. Spectra of peptides identified as acetylated were manually inspected for the presence of acetylation marker ions^53,54^.

### Phylogenetic conservation analyses

Amino acid sequences from Vav family proteins were obtained from the UniProt database (National Center for Biotechnology Information, Bethesda, MD) and aligned using the multiple sequence alignment by log-expectation algorithm in the Jalview software^55-57^.

## 3D structures

3D structures were generated in PyMol and the Protein Data Bank-stored crystal structure files for the Vav1–Rac1 complex (3BJI), the Vav1 CH–Ac–DH–PH–ZF (3KY9), the Vav1 KRR–NSH3 (1K1Z), the Vav1 SH2 (2MC1), and the Vav1 CSH3 (2KBT) regions.

### Luciferase reporter assays

Exponentially growing Jurkat cells (2 x 10^7^ cells per condition) were electroporated (250V, 950 μF pulses) using a Gene Pulser II apparatus (Cat. number 165-2106, BioRad). In the case of JNK assays, the cells were electroporated with the firefly luciferase reporter pFR– Luc (5 μg), pFA2–c–Jun (2 μg), a vector constitutively expressing the Renilla luciferase (pRL–SV40, 10 ng), and 20 μg of the indicated expression vector. In the case of NF–AT assays, the cells were electroporated with 3 μg of a luciferase reporter plasmid containing three NFAT binding sites (pNFAT/luc), 10 ng of pRL–SV40, and the indicated experimental plasmids as above. When required, empty plasmids were included in the electroporations to maintain constant the total amount of DNA introduced in cells among all the experimental samples. After 36 hours in complete medium, the electroporated cells were either left untreated or stimulated with the indicated antibodies for seven hours. Cells were then washed in serum-free media, lysed in Passive Lysis Buffer (Cat. number E1960; Promega), and the luciferase activity obtained in each condition recorded using the Dual-Luciferase Reporter System (Cat. number E1960, Promega) according to the supplier’s recommendations in a Lumat LB 9507 luminometer (Berthold). The raw values obtained were normalized according to the activity of the *Renilla* luciferase recorded in each sample. Final values are represented as the fold-change of the normalized luciferase activity obtained when compared to the control sample. In all cases, the abundance of the ectopically expressed proteins under each experimental condition was verified by analyzing aliquots of the cell extracts used in the luciferase experiments by immunoblot.

### Image processing

All images and figures were assembled and processed for final presentation with Canvas Draw 2 for Mac software (version 2.0, Canvas X).

### Statistical analyses

All the statistical analyses were carried out using GraphPad Prism software (version 6.0 and 8.0). The number of replicates and the statistical test used in each case are indicated in the figure legends. In all cases, the *P* values have been depicted using the * (when < 0.05), ** (when < 0.01), and *** (when < 0.001) notation.

## ACKNOWLEDGEMENTS

We thank M. Blázquez, A.L. Prieto, N. Ibarrola, and R. Dégano for technical assistance. X.R.B. is supported by grants from the Castilla-León Government (CLC-2017-01), the Spanish Ministry of Science, Innovation and Universities (MSIU) (RTI2018-096481-B-I00), and the Spanish Association against Cancer (GC16173472GARC). X.R.B.’s institution is supported by the Programa de Apoyo a Planes Estratégicos de Investigación de Estructuras de Investigación de Excelencia of the Ministry of Education of the Castilla-León Government (CLC-2017-01). S.R-F. and L.F.L-M. contracts have been supported by funding from the MISIU (S.R-F., BES-2013-063573), the Spanish Ministry of Education, Culture and Sports (L.F.L-M., FPU13/02923), and the CLC-2017-01 grant (S.R-F. and L.F.L-M.). L.F.N. is supported by the Salamanca local section of the Spanish Association for Cancer Research. Both Spanish and Castilla-León government-associated funding is partially supported by the European Regional Development Fund.

## AUTHOR CONTRIBUTIONS

S.R-F. designed and carried out most of experiments, analyzed the data, and performed the artwork and manuscript writing. L.F.N. and L.F.L-M. helped carried out experimental work. X.R.B. conceived the work, analyzed data, and carried out the final editing of the figures and manuscript.

---

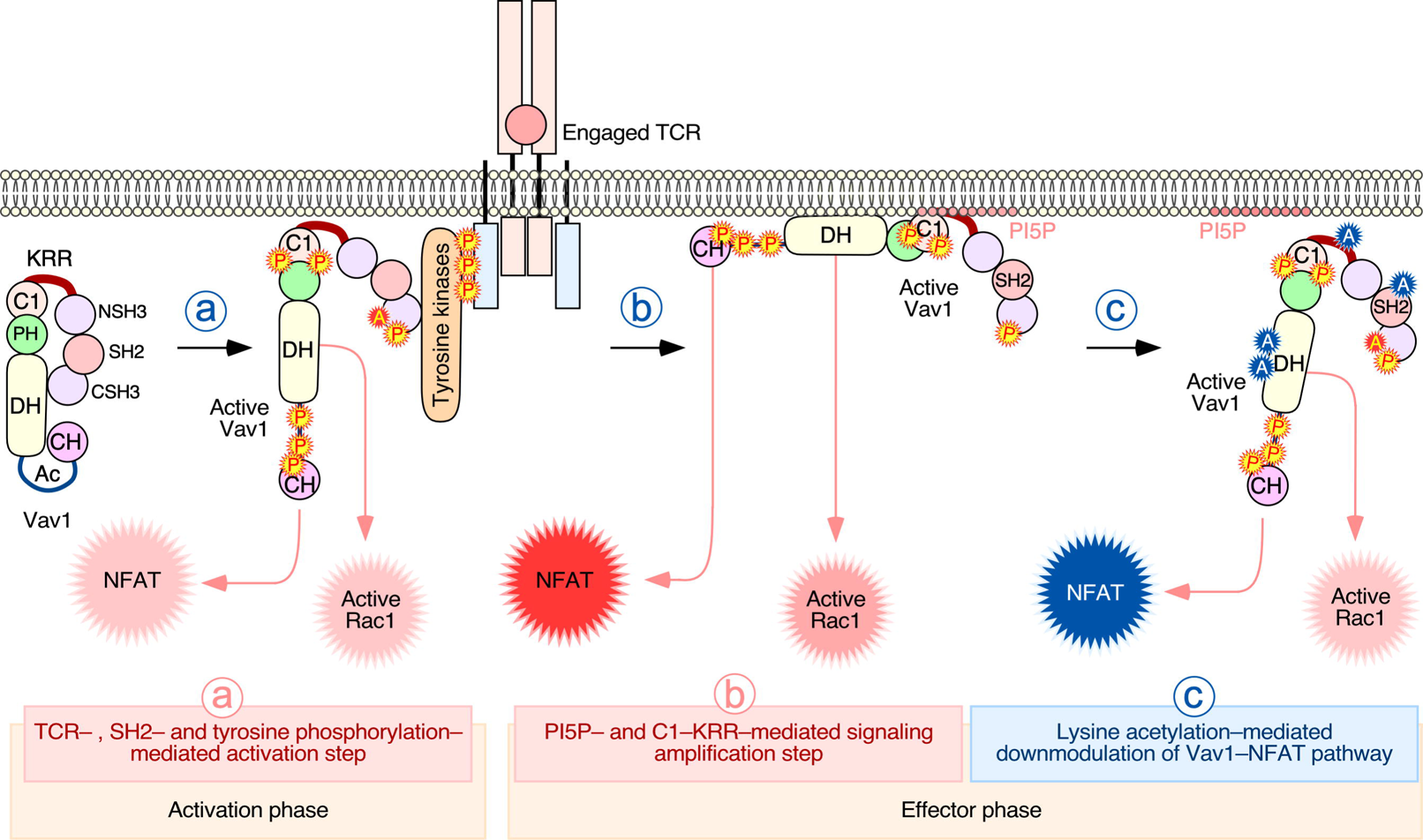

