## Supplemental Information for "Lysine Acetylation Reshapes The Downstream Signaling Landscape of Vav1 in Lymphocytes"

by

Sonia Rodríguez–Fdez, Lucía F. Nevado, L. Francisco Lorenzo–Martín and Xosé R. Bustelo*

This PDF file includes:

**(1)** Supplemental Figures S1 to S4 and legends (pages 2 to 7)

**(2)** Supplemental Table S1 (page 8)

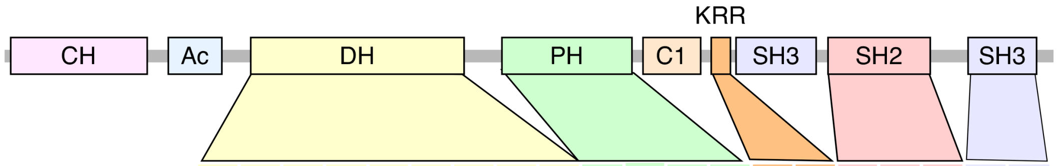


| Species | | K194 | K222 | K252 | K292 | K307 | K335 | K345 | K353 | K374 | K429 | K435 | K444 | K487 | K587 | K588 | K716 | K732 | K733 | K782 | K815 |
|---|---|---|---|---|---|---|---|---|---|---|---|---|---|---|---|---|---|---|---|---|---|
| *Capsaspora owczarzaki* | | * | * | – | – | – | * | * | – | * | – | – | * | * | – | – | * | – | – | * | – |
| *Salpingoeca rosetta* | | – | – | – | – | – | – | * | – | * | – | * | * | * | – | – | – | – | * | – | – |
| *Amph. queenslandica* | | * | – | – | – | – | – | * | – | * | * | – | * | * | – | – | * | – | – | – | – |
| *Trichoplax sp. H2* | | * | – | – | – | – | – | * | – | * | – | * | * | * | – | – | * | – | – | * | – |
| *Hydra vulgaris* | | * | * | – | – | * | * | * | – | * | – | * | * | * | – | – | * | – | * | * | – |
| *Clonorchis sinensis* | | – | – | – | – | – | * | * | – | * | * | * | – | * | – | – | – | – | – | * | – |
| *Caenorhabditis elegans* | | * | – | – | * | – | * | * | – | – | * | * | * | * | – | – | * | – | – | – | – |
| *Lottia gigantea* | | * | * | – | – | * | * | * | * | * | * | * | * | * | – | – | – | – | * | – | – |
| *Helobdella robusta* | | * | * | * | – | – | * | * | – | – | * | * | * | * | – | – | * | – | – | * | – |
| *Drosophila melanogaster* | | – | – | – | – | – | * | * | – | * | * | * | – | * | – | – | * | – | – | – | – |
| *Daphnia magna* | | * | – | – | – | * | * | * | – | – | _ | * | * | * | – | – | * | – | – | – | – |
| *Branchiostoma floridae* | | * | – | – | – | * | * | * | * | * | * | * | * | * | – | – | * | – | * | – | – |
| *Ciona intestinalis* | | * | – | – | – | – | * | * | * | * | _ | * | * | * | * | * | * | – | * | – | – |
| *Homo sapiens* | Vav2 | * | – | – | – | * | * | – | – | * | * | * | * | * | – | – | * | – | * | – | – |
| | Vav3 | – | – | – | – | * | * | * | * | * | – | * | * | * | – | * | * | – | – | * | – |

### FIGURE S1. Phylogenetic conservation of the indicated Vav1 acetylation sites

Black asterisks indicate the presence of a lysine residue in the primary sequence of the indicated Vav family protein that is conserved in mouse Vav1. Lysine residues identified as acetylation sites in other Vav family proteins using high-throughput proteomics analyses are indicated in red.

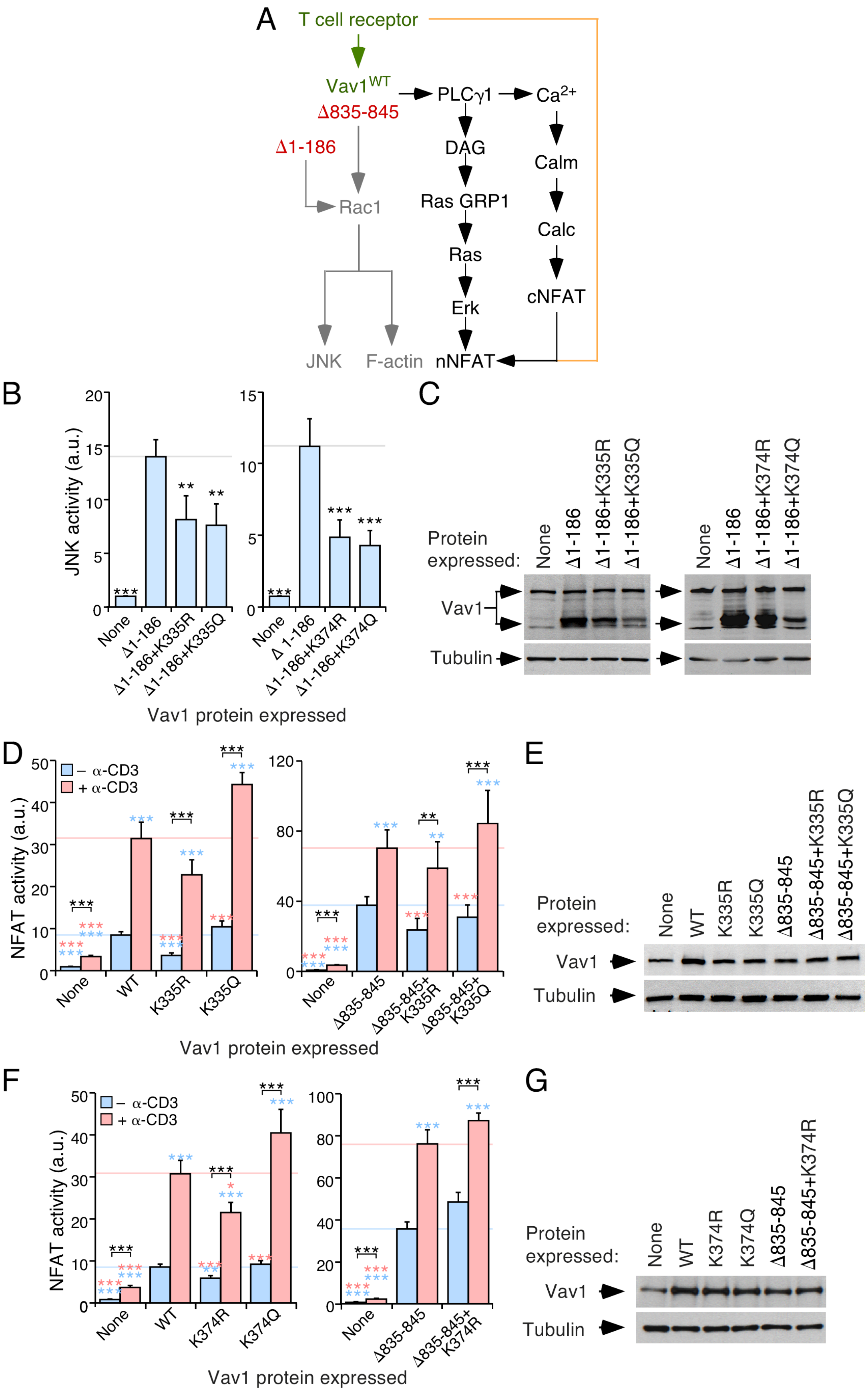
A
T cell receptor
Vav1WT
Δ835-845
Δ1-186
PLCγ1
Ca2+
DAG
Calm
Rac1
Ras GRP1
Calc
Ras
cNFAT
Erk
JNK
F-actin
nNFAT
B
JNK activity (a.u.)
None
Δ1-186
Δ1-186+K335R
Δ1-186+K335Q
Δ1-186+K374R
Δ1-186+K374Q
Vav1 protein expressed
C
Protein expressed:
Vav1
Tubulin
D
– α-CD3
+ α-CD3
NFAT activity (a.u.)
WT
K335R
K335Q
Δ835-845
Δ835-845+K335R
Δ835-845+K335Q
Vav1 protein expressed
E
F
K374R
K374Q
Δ835-845+K374R
G

### FIGURE S2. Mutations targeting $K^{335}$ and $K^{374}$ impair Vav1 catalysis–dependent pathways

**(A)** Schematic representation of the Vav1–dependent pathways that have been characterized in T cells. The Rac1–dependent pathway directly stimulated by the catalytic activity of Vav1 is shown in gray. The Vav1 adaptor–like pathway involved in the activation of NFAT is shown in black. Synergies established with parallel, TCR/CD3–triggered pathways are shown in brown. The WT and constitutively active versions of Vav1 used in this study are shown in green and red, respectively. DAG, diacylglycerol; GRP, GDP releasing factor; Calm, calmodulin; Calc, calcineurin; cNFAT, cytosolic NFAT; nNFAT, nuclear NFAT.

**(B)** Levels of stimulation of JNK triggered by the indicated Vav1 proteins in nonstimulated Jurkat cells. Data represent the mean ± SEM. **, $P < 0.01$; ***, $P < 0.001$ using the Mann–Whitney U test of indicated experimental values compared to $\text{Vav1}^{\Delta 1-186}$–expressing cells ($n = 3$ independent experiments, each performed in triplicate).

**(C)** Representative immunoblots showing the abundance of the ectopic Vav1 proteins and endogenous tubulin α in the experiments shown in B.

**(D,F)** Activation levels of NFAT elicited by the indicated Vav1 proteins in nonstimulated and TCR–stimulated Jurkat cells. Data represent the mean ± SEM. Statistical values were obtained using the Mann–Whitney U test. Blue and salmon asterisks indicate the significance level compared with nonstimulated and TCR–stimulated $\text{Vav1}^{\text{WT}}$–expressing cells (left panel) or $\text{Vav1}^{\Delta 835-845}$–expressing cells (right panel), respectively ($n = 3$ independent experiments, each performed in triplicate).

**(E,G)** Representative immunoblots showing the abundance of the ectopic Vav1 proteins and endogenous tubulin α in the experiments shown in D (E) and F (G).

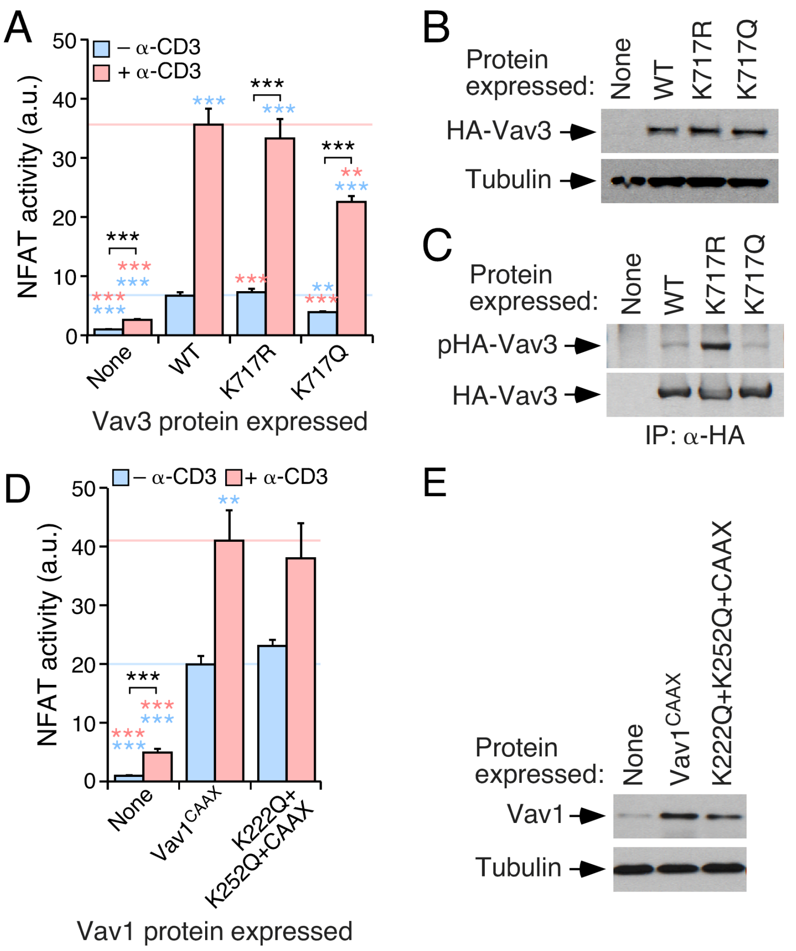


**FIGURE S3. The defects associated with the acetylation of Lys$^{716}$ are Vav1–specific**

**(A)** Activation levels of NFAT triggered by indicated HA–tagged Vav3 proteins in nonstimulated and TCR–stimulated Jurkat cells. Data represent the mean ± SEM. Statistical values were obtained using the Mann–Whitney U test. Blue and salmon asterisks indicate the significance level compared with nonstimulated and TCR–stimulated HA–Vav3$^{WT}$–expressing cells, respectively. Black asterisks refer to the *P* values obtained between the indicated experimental pairs (in brackets). *n* = 3 independent experiments, each performed in triplicate.

**(B)** Representative immunoblots showing the abundance of the ectopic HA–Vav3 proteins and endogenous tubulin α in the experiment shown in A.

**(C)** Tyrosine phosphorylation levels of the indicated HA–Vav3 proteins (top) immunoprecipitated from exponentially growing COS1 cells. Similar results were obtained in 3 independent experiments.

**(D)** Activation levels of NFAT induced by the indicated Vav1 proteins in in nonstimulated and TCR–stimulated Jurkat cells. Data represent the mean ± SEM. Statistical values were obtained using the Mann–Whitney U test. Blue and salmon asterisks indicate the significance level compared with nonstimulated and TCR–stimulated Vav1$^{CAAX}$–expressing cells, respectively (*n* = 3 independent experiments, each performed in triplicate).

**(E)** Representative immunoblots showing the abundance of the ectopic Vav1 proteins and endogenous tubulin $\alpha$ in the experiments shown in D.

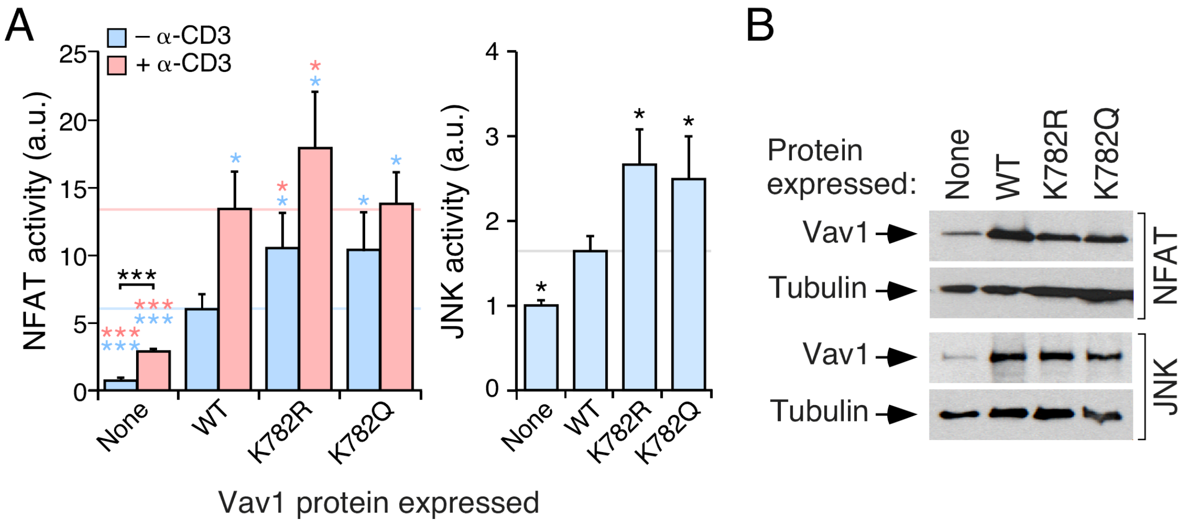


**FIGURE S4. Mutations targeting the Lys[782] promote the stimulation of Vav1 downstream signaling**

**(A)** Activation of NFAT (left panel) and JNK (right panel) by indicated Vav1 proteins in nonstimulated and TCR–stimulated Jurkat cells. Data represent the mean ± SEM. Statistical values were obtained using the Wilcoxon matched–pairs signed rank test. Blue and salmon asterisks indicate the significance level compared with nonstimulated and TCR–stimulated Vav1$^{WT}$–expressing cells, respectively. $n$ = 3 (NFAT assays) and 4 (JNK assays) independent experiments, each performed in triplicate.

**(B)** Representative immunoblots showing the abundance of the ectopic Vav1 proteins and endogenous tubulin α in the experiments shown in A.

### TABLE S1. Sequence of oligonucleotides used in this study

| Mutant | DNA sequence of primer | |
|---|---|---|
| Vav1$^{K222R}$ | F* | 5'– CAGCAGCACTTCATGAGGCCTCTGCAGCTATTC –3' |
| | R | 5'– GAATCGCTGCAGAGGCCTCATGAAGTGCTGCTG–3' |
| Vav1$^{K222Q}$ | F | 5'– CAGCACTTCATGCAGCCTCTGCAGC –3' |
| | R | 5'– GCTGCAGAGGCTGCATGAAGTGCTG –3' |
| Vav1$^{K252R}$ | F | 5'– GCATACCCACTTCTTAAGGGAACTGAAGGATGCCC –3' |
| | R | 5'– GGGCATCCTTCAGTTCCCTTAAGAAGTGGGTATGC–3' |
| Vav1$^{K252Q}$ | F | 5'– GCATACCCACTTCTTACAGGAACTGAAGGATGC –3' |
| | R | 5'– GCATCCTTCAGTTCCTGTAAGAAGTGGGTATGC –3' |
| Vav1$^{K335R}$ | F | 5'– ATGCAGCGGGTGCTGAGGTACCACCTCCTTCTC –3' |
| | R | 5'– GAGAAGGAGGTGGTACCTCAGCACCCGCTGCAT–3' |
| Vav1$^{K335Q}$ | F | 5'– CCTATGCAGCGGGTGCTGCAGTACCACCTCCTTCTCC –3' |
| | R | 5'– GGAGAAGGAGGTGGTACTGCAGCACCCGCTGCATAGG –3' |
| Vav1$^{K374R}$ | F | 5'– TGCGTGAACGAGGTCAGGAGGGACAATGAAACC–3' |
| | R | 5'– GGTTTCATTGTCCCTCCTGACCTCGTTCACGCA –3' |
| Vav1$^{K374Q}$ | F | 5'– GTGCGTGAACGAGGTCCAGAGGGACAATGAAAC –3' |
| | R | 5'– GTTTCATTGTCCCTCTGGACCTCGTTCACGCAC –3' |
| Vav1$^{K587R}$ | F | 5'– GGCCCAGGACAGGAAAAGGAATG –3' |
| | R | 5'– CATTCCTTTTCCTGTCCTGGGCC –3' |
| Vav1$^{K587Q}$ | F | 5'– GGGCCCAGGACCAGAAAAGGAATG –3' |
| | R | 5'– CATTCCTTTTCTGGTCCTGGGCCC –3' |
| Vav1$^{K588R}$ | F | 5'– GGCCCAGGACAAGAGAAGGAATGAATTGG –3' |
| | R | 5'– CCAATTCATTCCTTCTCTTGTCCTGGGCC –3' |
| Vav1$^{K588Q}$ | F | 5'– GGGCCCAGGACAAGCAAAGGAATGAATTG –3' |
| | R | 5'– CAATTCATTCCTTTGCTTGTCCTGGGCCC –3' |
| Vav1$^{K716R}$ | F | 5'– CAGCATTAAGTATAACGTGGAGGTCAGGCATATTAAAATCATGACGTCAGA –3' |
| | R | 5'– CCTCTGACGTCATGATTTTAATATGCCTGACCTCCACGTTATACTTAATGC–3' |
| Vav1$^{K716Q}$ | F | 5'– CATTAAGTATAACGTGGAGGTCCAGCATATTAAAATCATGACGTC –3' |
| | R | 5'– GACGTCATGATTTTAATATGCTGGACCTCCACGTTATACTTAATG –3' |
| Vav1$^{K782R}$ | F | 5'– CAGCTGGAAGCACCAGGTATTTTGGCACTGC –3' |
| | R | 5'– GCAGTGCCAAAATACCTGGTGCTTCCAGCTG –3' |
| Vav1$^{K782Q}$ | F | 5'– CCAGCTGGAAGCACCCAGTATTTTGGCACTG –3' |
| | R | 5'– CAGTGCCAAAATACTGGGTGCTTCCAGCTGG –3' |
| Vav3$^{K717R}$ | F | 5'– GTACAATAATGAAGCAAGGCACATCAAGATTTTAAC –3' |
| | R | 5'– GTTAAAATCTTGATGTGCCTTGCTTCATTATTGTAC –3' |
| Vav3$^{K717Q}$ | F | 5'– CAATAATGAAGCACAGCACATCAAG –3' |
| | R | 5'– CTTGATGTGCTGTGCTTCATTATTG –3' |

*F, forward primer; R, reverse primer. Nucleotides that have been replaced in the WT *Vav1* coding sequence to incorporate the indicated point mutations are shown in red.
